# Redesign selective protein binders using contrastive decoding

**DOI:** 10.64898/2026.05.09.722041

**Authors:** Ziwei Xie, Jinbo Xu

**Affiliations:** Toyota Technological Institute at Chicago, Chicago, IL 60637, United States

## Abstract

**Motivation:** Fixed-backbone sequence design methods such as ProteinMPNN operate on backbone coordinates alone and cannot represent target side-chains at the binding interface. Their decoding algorithm also lacks a mechanism to balance binding affinity and folding stability or to improve selectivity against structurally similar off-targets. These gaps limit the computational design of protein binders with high affinity and specificity.

**Results:** We present RedNet, a multiscale graph neural network that encodes side-chain information of the binding target. We further develop a contrastive decoding algorithm, motivated by the thermodynamic decomposition of binding free energy, that addresses two objectives: (1) balancing binding affinity and folding stability, and (2) improving selectivity against structurally similar off-targets. RedNet reaches 43% native sequence recovery on heterodimers, compared with 37% for ProteinMPNN and 33% for ESM-IF. With contrastive decoding, it matches native-sequence co-folding success (68%) on high-confidence AlphaFold3 targets, exceeding ProteinMPNN (59%) and ESM-IF (61%). On a new benchmark of structurally similar on-/off-target pairs, RedNet with contrastive decoding reaches 64.8% energetic selectivity, ahead of PiFold (55.6%), ProteinMPNN (53.7%), and ESM-IF (53.7%).

**Availability:** Source code and datasets are released at https://github.com/zw2x/rednet_public.

## 1 Introduction

Designing protein binders that engage a chosen target with high affinity and specificity is a central problem in protein engineering, with applications spanning therapeutics and basic research [16, 15, 4]. Recent deep learning methods have expanded design capabilities, but producing binders that are simultaneously potent and selective, without extensive experimental screening, remains difficult [24, 20].

Structure-based binder design, or one-sided interface design [16], typically proceeds in three stages [24, 20, 22]: (1) generating a backbone or all-atom structure conditioned on the target, (2) redesigning sequences based on the fixed structures to optimize binding and other properties, and (3) filtering and ranking candidates with machine-learning and physics-based scores. Recent progress in each stage has been substantial, but current fixed-backbone sequence design methods do not address the particular needs of binder design: affinity optimization without destabilizing the fold, discrimination against off-targets, and accurate modeling of the side-chain conformations that define affinity and specificity.

This paper addresses the sequence redesign stage. ProteinMPNN is the most widely adopted fixed-backbone model; it is efficient and has robust performance on monomers with idealized scaffolds. Two gaps limit it for binder sequence redesign. First, it sees only backbone atoms and cannot capture side-chain conformations at the target, which are central to binding affinity and specificity. Second, its decoding algorithm has no explicit mechanism to jointly optimize protein complex binding affinity while maintaining binder folding stability, or to favor on-target interactions over structurally similar off-target interactions.

To close these gaps, we introduce RedNet, a multiscale graph neural network that incorporates backbone geometry and target side-chain information, enabling more accurate modeling of binding interfaces. We further introduce a contrastive decoding algorithm derived from the thermodynamic decomposition of binding free energy, which improves binding while preserving folding stability and widens the free energy gap between on-target and off-target interactions.

We evaluate RedNet on three tasks. For native sequence recovery, RedNet reaches 43% on heterodimers, ahead of ProteinMPNN (37%) and ESM-IF (33%). For AlphaFold3 co-folding success on high-confidence targets, RedNet with contrastive decoding (RedNet-CD) matches native sequences at 68% success, above ProteinMPNN (59%), ESM-IF (61%), and PiFold (64%). Rosetta energetics show that RedNet-CD and an ensemble variant (RedNet-Ens) produce binding scores, hydrogen bonding, and surface hydrophobicity matching or exceeding those of native interfaces. On a new selective-binder benchmark from the PDB, RedNet-CD attains 64.81% energetic selectivity at the base threshold, nearly doubling that of RedNet without contrastive decoding (33.33%) and leading all baselines.

### Related work

Protein design methods fall broadly into physics-based and deep learning approaches. Physics-based methods combine energy functions with stochastic (e.g., Rosetta [2, 21]) or deterministic (e.g., OSPREY [10]) search, but scale poorly and often require extensive experimental screening. Deep learning fixed-backbone methods, both autoregressive, such as Structured Transformer [14] and ProteinMPNN [5], and non-autoregressive, such as PiFold [7] and Frame2Seq [1], have substantially improved efficiency and experimental success rates [5, 1]. For one-sided interface design, pipelines such as RFdiffusion [22] combined with ProteinMPNN, and BindCraft [20], have produced experimentally validated binders, though success rates vary widely across targets [24]. Multistate design for binding specificity has been addressed via Rosetta MSD [13], deep learning extensions [11, 9], and experimental heuristics [25], but whether these approaches generalize to systematically tuning specificity remains unclear. See the supplement for a detailed discussion.

## 2 Methods

### 2.1 Protein graph representation

We use multi-scale graph representations to model the backbone and side-chain geometry of protein complexes.

#### Backbone representation for complex structures

We represent the complex as a residue-level graph *G* = (*V, E*), where nodes *v*_*i*_ ∈*V* correspond to residues and edges *e*_*ij*_ ∈*E* connect spatially proximal residues within a distance cutoff. Local frames are derived from backbone atom coordinates (N, C*α*, C, O) and edges are built by *k*-nearest neighbors on C*α* distances, following GLINTER [23].

#### Side-chain representation for target chains

For target chains with known sequence, we build an all-atom graph *G*_atom_ = (*V*_atom_, *E*_atom_) to capture side-chain interactions. Each node *v*_*a*_ ∈*V*_atom_ represents a heavy atom, with features encoding its local chemical environment. Edges connect atoms within a distance cutoff (e.g., 10 Å) and encode pairwise distances and additional features; see full list of node and edge features in the supplement.

### 2.2 Network architectures

#### 2.2.1 Overview

Figure 1 shows the overall architecture. Graph neural networks encode the protein graphs; a causal transformer decodes amino acids at each timestep. The building blocks below are variants of standard GAT and attention layers tailored to causal sequence decoding over protein graphs.

**Figure 1:**
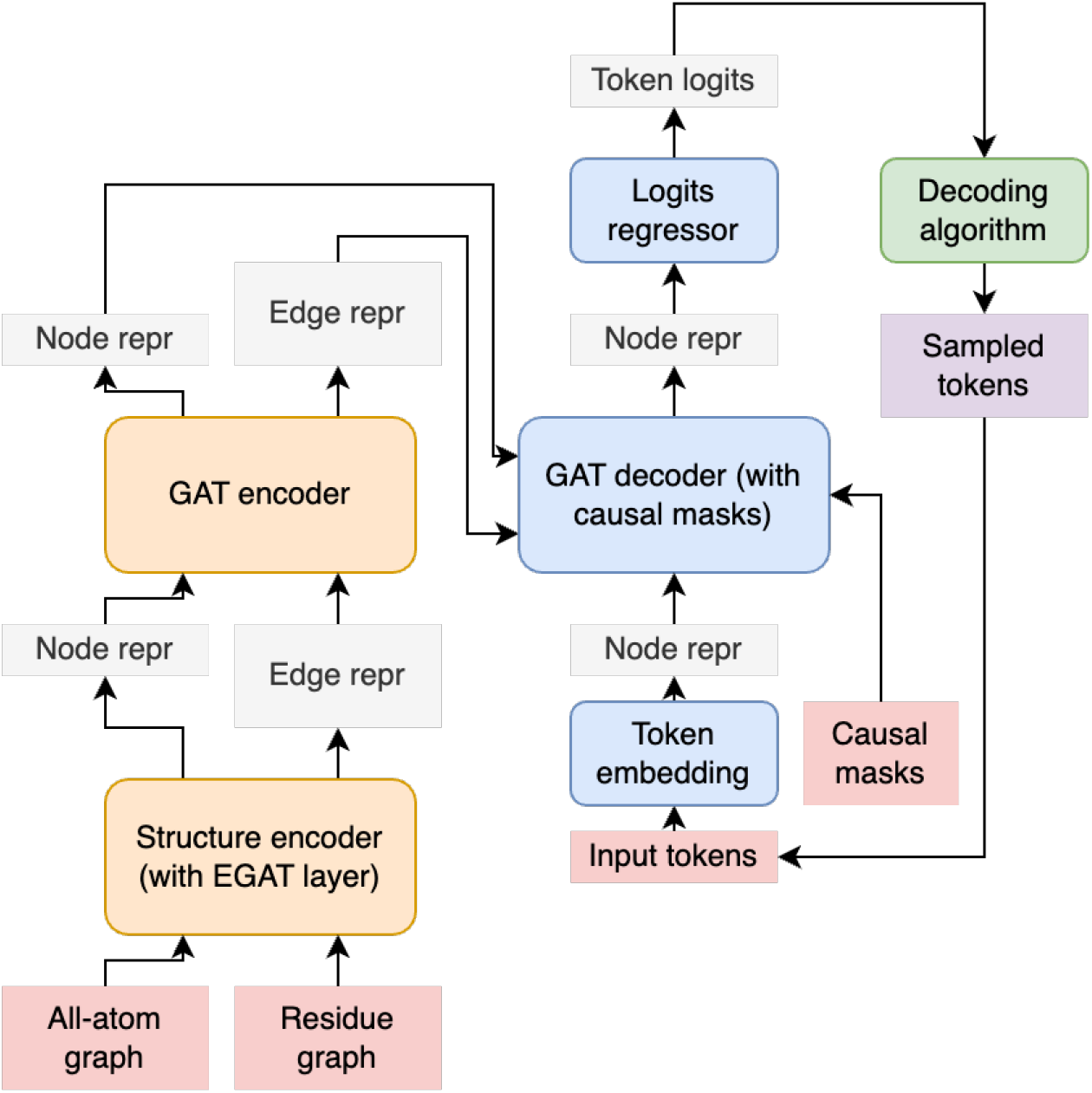
Overview of the RedNet architecture. Graph neural networks encode protein structure into node and edge representations, which are then decoded by a causal graph network to autoregressively predict amino acid sequences.

#### Graph attention network

Our GAT variant (Supplementary Algorithm S1) combines a mean-pooled global branch with a GATv2-style attention branch [3], both modulated by zero-initialized sigmoid gates for stable training. Causal masks over the sampled decoding order restrict each residue to previously decoded neighbors. For target side-chain modeling we add an SE(3)-invariant all-atom GAT; see supplement for the full architecture and pseudocode.

#### Loss

We train with a cross-entropy loss over amino acid tokens [14] and add an auxiliary edgewise cross-entropy regularizer; full definitions are in the supplement.

### 2.3 Contrastive decoding

Contrastive decoding [17, 18] modifies the predicted logits at each timestep to amplify features that distinguish the on-target bound structure from an off-target one. Here, a bound structure consists of the backbone coordinates of the complex and the side-chain coordinates of the target chain. At timestep *t*, the modified logits are

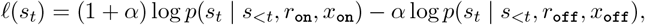

where *s*_*t*_ is the binder residue at position *t, s*_<*t*_ denotes the previously sampled binder residues, *r*_on_, *r*_off_ are the on- and off-target sequences, *x*_on_, *x*_off_ the corresponding bound structures, and *α* ≥0 controls the contrastive strength. *α* = 0 recovers standard decoding on the on-target distribution; as *α* grows, residue choices that are also favorable under the off-target are penalized.

To prevent the contrastive term from selecting low-probability tokens, we restrict the candidate set at each timestep to tokens with sufficient probability under the on-target:

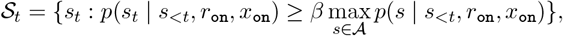

where 𝒜 is the amino acid alphabet and *β* ∈[0, 1] is a truncation threshold. We then sample from *p*_*t*_(*s*) = softmax_𝒮*t*_ (*𝓁*(*s*)).

The same framework enables sampling sequences that optimize the binding free energy Δ*G*. The free-energy decomposition [8] is

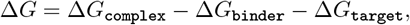

with Δ*G*_complex_ the free energy of the bound complex and Δ*G*_binder_, Δ*G*_target_ the free energies of the unbound binder and target. Folding free energy correlates with the log-likelihood of finetuned fixed-backbone design models [6], so

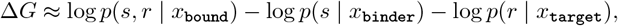

where *s, r* are the binder and target sequences and *x*_bound_, *x*_binder_,*x*_target_ the corresponding structures. Since *r* is fixed during binder design, dropping *s*-independent terms gives

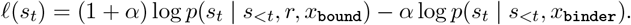

This selects residues favorable in the bound state but unfavorable in the unbound state, approximating the thermodynamic preference for complex formation. The same candidate-set truncation and sampling procedure apply. The same bound/unbound contrast also yields a family of scoring metrics for zero-shot binding affinity prediction; see Supplementary Section S1 for definitions and SKEMPI v2.0 results.

### 2.4 Datasets and benchmarks

#### Training and validation

We train on PDB structures released before 2023-01-01, filtered to X-ray/EM entries with resolution ≤ 5 Å, ≤ 20 polymer chains, and polypeptide(L) chains of 10–5000 residues with < 10% unknown residues. Structures released between 2022-05-01 and 2022-12-31 form the validation set, with redundancy removed as for the test set.

#### Low-homology test set

We select structures released 2023-01-01 to 2023-12-31 and retain only chains with MMseqs2 e-value > 1 against training sequences, stricter than the common 30% identity cutoff. Remaining sequences are clustered at 40% identity, and we sample 300 monomers, 150 homodimers, and 150 heterodimers from cluster representatives. Heterodimer co-folding is evaluated on the 107 heterodimers with ≤ 500 total residues.

#### Selective binder benchmark

We assemble a selective binder benchmark from PDB heterodimers released before 2025-04-14. For each binder cluster, one interacting heterodimer is the on-target; others in the cluster become candidate off-targets. We keep candidates whose binder chains align with coverage ≥90%, identity ≥90%, and RMSD ≤ 2.5 Å to the on-target binder, and whose target chains are not identical. Interface similarity is measured by the Jaccard index over inter-chain C*α*–C*α* contact pairs (≤10 Å); pairs with Jaccard ≥ 0.9 are discarded. For each cluster we take the off-target with lowest Jaccard, yielding 656 non-redundant pairs, from which we uniformly sample 180 for evaluation. Full curation details are in the supplement.

#### 2.4.1 Benchmarking methods

We compare against widely adopted fixed-backbone sequence design methods: ESM-IF [12], ProteinMPNN [5], and PiFold [7].

## 3 Results

### 3.1 All-atom graph transformer improves sequence recovery on monomeric and dimeric structures

We benchmark RedNet against ESM-IF, ProteinMPNN, and PiFold on native sequence recovery (NSR), wild-type log-likelihood (LL), and perplexity (PPL) across monomers, homodimers, and heterodimers (Table 1). ESM-IF and PiFold are trained without noise augmentation; ProteinMPNN and RedNet are compared across *σ* ∈ {0, 0.02, 0.1, 0.2, 0.3} to probe robustness to backbone coordinate perturbation.

**Table 1:** Performance comparison on monomers, homodimers, and heterodimers. *σ*: backbone coordinate noise level. NSR: Native Sequence Recovery. LL: Log-Likelihood. PPL: Perplexity. For RedNet and ProteinMPNN, we test performance at different noise levels *σ* ∈{0.02, 0.1, 0.2, 0.3}. ESM-IF and PiFold are tested at *σ* = 0.

| Model | $\sigma$ | Monomer | | | Homodimer | | | Heterodimer | | |
| --- | --- | --- | --- | --- | --- | --- | --- | --- | --- | --- |
|  |  | NSR | LL | PPL | NSR | LL | PPL | NSR | LL | PPL |
| ESM-IF | 0 | 0.38 | -1.85 | **7.33** | 0.43 | -1.35 | 4.29 | 0.33 | -2.21 | 13.39 |
| PiFold | 0 | 0.40 | -1.92 | 9.65 | 0.45 | -1.73 | 6.10 | 0.35 | -2.08 | 9.16 |
| RedNet (ours) | 0 | **0.43** | **-1.74** | 9.15 | **0.49** | **-1.28** | **3.86** | **0.43** | **-1.76** | **6.58** |
| ProteinMPNN | 0.02 | 0.36 | **-1.91** | **7.60** | 0.42 | -1.55 | 5.12 | 0.37 | -2.00 | 8.32 |
| RedNet (ours) | 0.02 | **0.37** | -1.92 | 8.30 | **0.43** | **-1.44** | **4.64** | **0.39** | **-1.91** | **7.51** |
| ProteinMPNN | 0.1 | **0.33** | -2.02 | **8.36** | **0.40** | -1.64 | 5.58 | 0.34 | -2.11 | 9.25 |
| RedNet (ours) | 0.1 | **0.33** | **-2.01** | 8.68 | **0.40** | **-1.53** | **5.07** | **0.35** | **-2.04** | **8.67** |
| ProteinMPNN | 0.2 | 0.31 | -2.08 | **8.84** | 0.37 | -1.70 | 5.90 | 0.33 | -2.14 | 9.46 |
| RedNet (ours) | 0.2 | **0.32** | **-2.06** | 8.91 | **0.38** | **-1.60** | **5.38** | **0.34** | **-2.08** | **8.84** |
| ProteinMPNN | 0.3 | **0.30** | -2.12 | **9.18** | **0.36** | -1.76 | 6.28 | 0.31 | -2.21 | 10.12 |
| RedNet (ours) | 0.3 | **0.30** | **-2.11** | 9.23 | **0.36** | **-1.63** | **5.54** | **0.32** | **-2.13** | **9.27** |

#### Monomers and homodimers

RedNet beats every baseline at low noise. On monomers it reaches 0.43 NSR against 0.38 (ESM-IF) and 0.40 (PiFold); on homodimers, 0.49 against 0.43 and 0.45. Perplexity and log-likelihood follow the same ordering. At *σ* ≥0.1, RedNet’s NSR matches ProteinMPNN’s, but it gives lower perplexity and higher log-likelihood.

#### Heterodimers

The gap is largest on heterodimeric interfaces, which matter most for binder design. At *σ* = 0, RedNet reaches NSR 0.43 against 0.33 (ESM-IF), 0.35 (PiFold), and 0.37 (ProteinMPNN, *σ* = 0.02). Perplexity separates the methods more sharply: RedNet 6.58 vs. ESM-IF 13.39, PiFold 9.16, ProteinMPNN 8.32. Log-likelihood follows the same ordering. We study the effects of different architectural components on sequence recovery at *σ* = 0.02, as shown in Supplementary S2. Using a GAT architecture with side-chain information improves heterodimer results (0.39 vs. 0.37).

#### Robustness to coordinate noise

RedNet’s heterodimer NSR drops from 0.43 (*σ* = 0) to 0.32 (*σ* = 0.3); ProteinMPNN’s drops from 0.37 (*σ* = 0.02) to 0.31. The NSR gap narrows as noise grows, but RedNet’s perplexity stays lower throughout (8.84 vs. 9.46 at *σ* = 0.2; 9.27 vs. 10.12 at *σ* = 0.3). The narrowing NSR gap likely reflects RedNet’s use of side-chain geometry, which degrades with backbone perturbation more than backbone-only features do.

### 3.2 Contrastive decoding improves co-folding success of designed binders

In binder design applications [20], AlphaFold3 confidence metrics (pLDDT, ipTM, pTM) are used to filter designs and have been shown to guide wet-lab selection. We withhold MSAs to prevent confidence inflation and rely on target chain templates, and use BindCraft’s default thresholds as success criteria.

We evaluate on all 107 heterodimeric targets (Table 2) and on a 44-target high-confidence subset where native-sequence AF3 predictions exceed pLDDT 70 (Table 3).

**Table 2:**
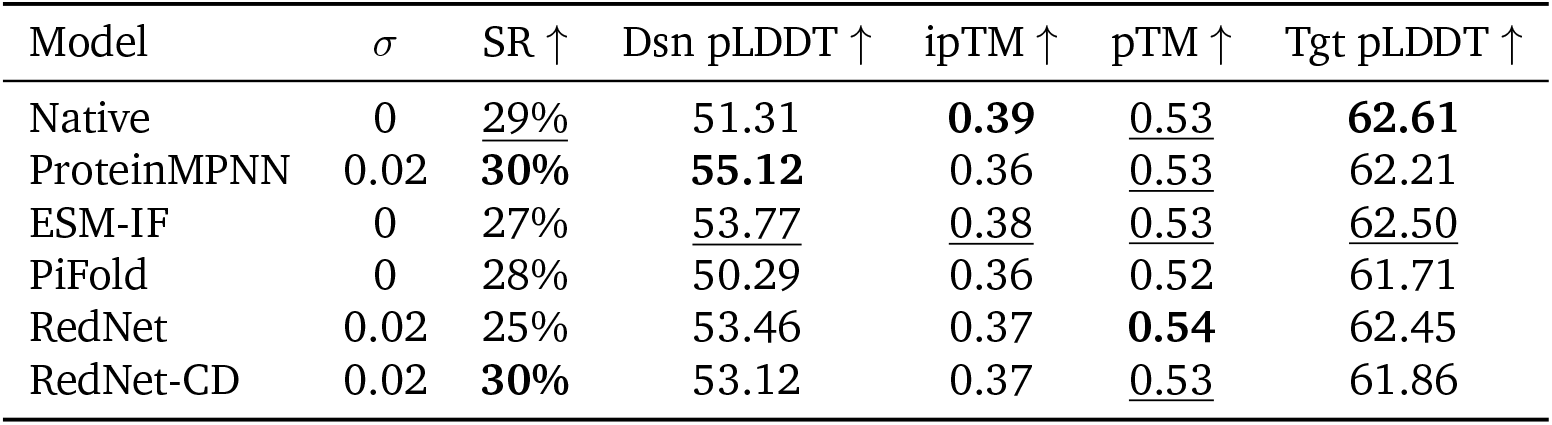
Heterodimer co-folding results on all 107 targets. *σ*: backbone coordinate noise level (Å). SR: Success Rate (0–100%), defined as pTM > 0.55, ipTM > 0.5, and Dsn pLDDT > 80, following BindCraft. Dsn pLDDT: AlphaFold3 predicted LDDT for the design chain (0–100). ipTM: interface predicted Template Modeling score (0–1). pTM: AlphaFold3 predicted TM-score of the complex (0–1). Tgt pLDDT: AlphaFold3 predicted LDDT for the target chain (0–100). RedNet-CD uses contrastive decoding with *α* = 1, *β* = 0.9. All models are sampled at temperature = 0.001. Higher is better for all metrics. **Bold**: best; <u>underline</u>: second best.

**Table 3:**
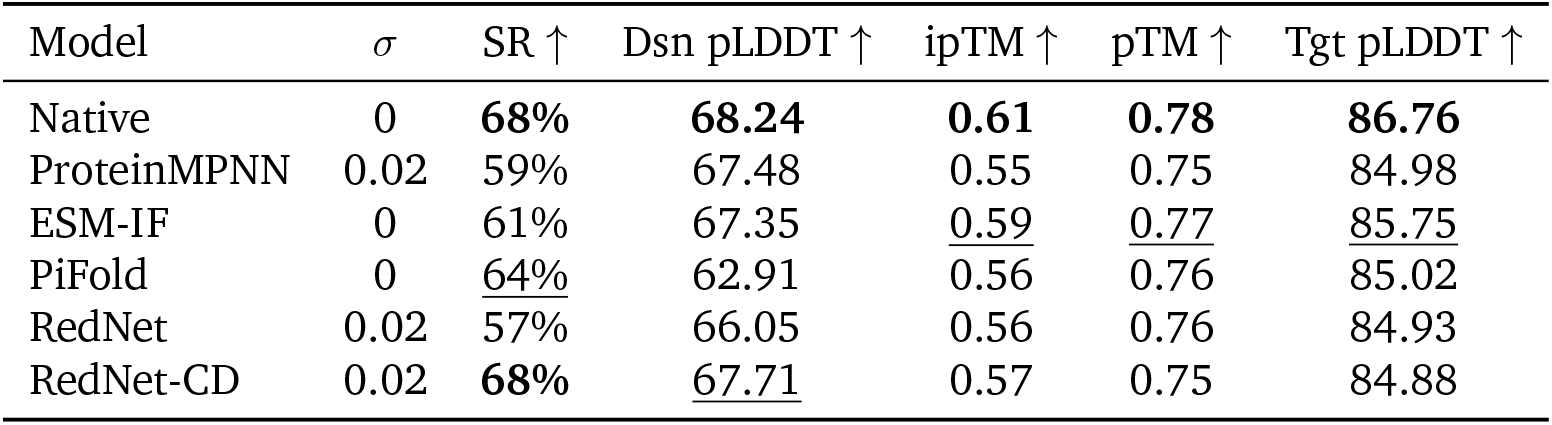
Co-folding performance on high-quality subset (44 targets). *σ*: backbone coordinate noise level (Å). SR: Success Rate (0–100%), defined as pTM > 0.55, ipTM > 0.5, and Dsn pLDDT > 80, following BindCraft. Dsn pLDDT: AlphaFold3 predicted LDDT for the design chain (0–100). ipTM: interface predicted Template Modeling score (0–1). pTM: AlphaFold3 predicted TM-score of the complex (0–1). Tgt pLDDT: AlphaFold3 predicted LDDT for the target chain (0–100). RedNet-CD uses contrastive decoding with *α* = 1, *β* = 0.9. All models are sampled at temperature = 0.001. Higher is better for all metrics. **Bold**: best; <u>underline</u>: second best.

#### All targets

On the full 107 heterodimers, RedNet-CD and ProteinMPNN both reach 30% success, matching or exceeding native sequences (Table 2). However, AlphaFold3 structure-prediction quality strongly limits the measurable success rate across all methods, as our factor analysis in Supplement S3 shows. We therefore focus on the high-confidence subset where design quality is the primary differentiator.

#### High-quality targets

On the 44 targets where native predictions are reliable, RedNet-CD matches native-sequence success at 68%, ahead of PiFold (64%), ESM-IF (61%), and Protein-MPNN (59%). Contrastive decoding raises RedNet’s own success from 57% to 68% on the same model, showing that the improvement comes from the decoding rule itself and not just the architecture. ProteinMPNN’s mean binder-chain pLDDT (67.48) is close to RedNet-CD’s (67.71), yet it trails on the composite success criterion, suggesting RedNet-CD clears the pLDDT > 80, pTM > 0.55, ipTM > 0.5 thresholds more often. We further study how different hyperparameters affect the co-folding success of the contrastive decoding algorithm, as shown in Supplement S2, and find *α* = 1 and *β* = 0.9 to be optimal at low temperature (*t* = 0.001).

#### Energetics

AlphaFold3 confidences correlate poorly with stability or affinity, so we additionally compute Rosetta energetics and biochemical properties following BindCraft [20], running three relaxation repeats per design and keeping the lowest-energy structure. We also report RedNet-Ens, which picks the better-scoring design between RedNet and RedNet-CD per target (Table 4; full hydrophobicity and hydrogen-bonding numbers in Supplementary Table S7). RedNet-Ens attains the most favorable binding score (−188.04 REU vs. −184.89 for ProteinMPNN and−172.42 for native), the best interface energy (−56.66 REU vs. native −53.35), and the largest buried surface area (1966 vs. native 1918 Å^2^). Contrastive decoding alone moves RedNet from −179.88 to −182.26 in binding score and from −52.49 to −54.47 in interface energy, with shape complementarity (0.66) and packing (0.54) comparable to baselines. ProteinMPNN has the tightest packing (Int Packstat 0.55) but the least favorable interface energy (−46.98), suggesting its designs over-optimize local packing at the expense of global interface energetics.

**Table 4:** Energetics and geometric properties of designed binders. Binding Score (REU): Rosetta binding score. Int SC (0–1): interface shape complementarity. Int Packstat (0–1): interface packing statistic. Int dG (REU): interface free energy change. Int dSASA (Å^2^): interface buried solvent-accessible surface area. REU: Rosetta Energy Units. Due to Rosetta relaxation failures, we analyze 91 of 107 heterodimers that are successfully relaxed for all methods. RedNet-Ens combines RedNet and RedNet-CD by selecting the design with the best binding score. **Bold**: best; <u>underline</u>: second best.

| Model | $\sigma$ | Binding Score ↓ | Int SC ↑ | Int Packstat ↑ | Int dG ↓ | Int dSASA ↑ |
| --- | --- | --- | --- | --- | --- | --- |
| Native | 0 | −172.42 | 0.65 | 0.53 | −53.35 | 1918.33 |
| ProteinMPNN | 0.02 | <u>−184.89</u> | <u>0.66</u> | **0.55** | −46.98 | 1682.12 |
| ESM-IF | 0 | −181.27 | 0.64 | 0.53 | −54.97 | <u>1947.75</u> |
| PiFold | 0 | −179.55 | **0.67** | 0.53 | <u>−55.19</u> | 1870.66 |
| RedNet | 0.02 | −179.88 | <u>0.66</u> | <u>0.54</u> | −52.49 | 1868.29 |
| RedNet-CD | 0.02 | −182.26 | <u>0.66</u> | <u>0.54</u> | −54.47 | 1894.69 |
| RedNet-Ens | 0.02 | **−188.04** | **0.67** | **0.55** | **−56.66** | **1965.96** |

#### Biochemical properties

Excessive surface hydrophobicity drives aggregation and off-target binding, so matching native levels is favorable. Native interfaces sit at 0.43 hydrophobicity; ProteinMPNN matches at 0.43, RedNet-CD and RedNet-Ens track closely at 0.44, and PiFold is highest at 0.47. Interface hydrogen bonds contribute to affinity, and buried polar atoms without partners destabilize the complex [19]. RedNet-Ens forms the most interface hydrogen bonds (7.31 per interface, 48.99% of interface residues; native 6.97, 46.15%) and the lowest fraction unsatisfied (14.27% vs. native 17.90%). RedNet-CD likewise exceeds native (7.01), while ProteinMPNN forms fewer (5.44), consistent with a more hydrophobic-packing-driven strategy. Contrastive decoding thus shifts RedNet toward hydrogen-bond-driven binding while keeping hydrophobicity near native levels. See details in the supplement.

### 3.3 Contrastive decoding improves binding specificity

We assemble a selective binder benchmark from PDB heterodimers. Each case pairs an on-target and an off-target receptor with similar backbones, so the binder must discriminate on side-chain features. We evaluate specificity two ways: Rosetta-based energetic selectivity (AlphaFold3 confidences are insensitive to mutational affinity changes) and AlphaFold3 co-folding selectivity via ipTM thresholds.

#### Energetic analysis

Following BindCraft [20], we define the Binder Score as Δ*G*_binding_ + Δ*G*_binder_ = Δ*G*_complex_ −Δ*G*_receptor_, which reflects both interface strength and binder stability in the bound state. A selective design should have a lower on-target than off-target Binder Score, so we measure selectivity by Score_on_ −Score_off_, with negative values favoring the on-target (Table 5). The benchmark contains 54 on-/off-target pairs filtered to Jaccard interface similarity ≤ 0.5. Native sequences, which were not optimized for selectivity between structurally similar receptors, succeed 50% of the time—consistent with random—confirming that the task requires active discrimination.

**Table 5:** Selectivity measured by Rosetta binding score difference (on-target −off-target). A negative value indicates the binder prefers the on-target. SR (Diff < X): percentage of cases where the on-target preference exceeds the energy gap threshold X. *σ*: backbone coordinate noise level (Å). Higher is better for all metrics. **Bold**: best; <u>underline</u>: second best.

| Model | $\sigma$ | SR (Diff < –10) ↑ | SR (Diff < –5) ↑ | SR (Diff < 0) ↑ |
| --- | --- | --- | --- | --- |
| Native | 0 | 25.93% | 33.33% | 50.00% |
| ProteinMPNN | 0.02 | 31.48% | 44.44% | 53.70% |
| ESM-IF | 0 | **35.19%** | 42.59% | 53.70% |
| PiFold | 0 | 27.78% | 40.74% | <u>55.56%</u> |
| RedNet | 0.02 | 22.22% | 27.78% | 33.33% |
| RedNet-CD | 0.02 | <u>33.33%</u> | **51.85%** | **64.81%** |

At Diff < 0, RedNet-CD leads at 64.81%, nearly double RedNet’s 33.33% and ahead of PiFold (55.56%), ProteinMPNN and ESM-IF (both 53.70%), and native (50%). At Diff < −5, RedNet-CD again leads at 51.85%, followed by ProteinMPNN (44.44%), ESM-IF (42.59%), PiFold (40.74%), native (33.33%), and RedNet (27.78%). Only at the strictest threshold (Diff < −10) does ESM-IF (35.19%) exceed RedNet-CD (33.33%). Contrastive decoding roughly doubles baseline RedNet’s selectivity at both looser thresholds. We further examine how hyperparameters affect the selective design success rate of the contrastive decoding algorithm, as shown in Supplementary S2. We find that *α* ∈ {1, 2} with *β* = 0.9 performs best across energy thresholds.

#### Co-folding analysis

We further examine the co-folding selectivity success rates in Table 6. RedNet-CD leads at 9.26RedNet-CD optimizes selectivity at the cost of some on-target binding. We note that the co-folding test can be more limited than the previous energetic analysis, because AlphaFold3 confidence is insensitive to the mutational changes that are important for distinguishing binding affinities of protein-protein interactions. However, the co-folding test still provides a standardized framework for comparing selectivity across methods, and we expect it to improve as co-folding models become more sensitive to mutational changes.

**Table 6:** Selectivity success measured by AlphaFold3 cofolding. *σ*: backbone coordinate noise level (Å). Selectivity: proportion where on-target ipTM > 0.55 and off-target ipTM < 0.55. On-Target: proportion where on-target ipTM > 0.55. Off-Target: proportion where off-target ipTM > 0.55. **Bold**: best; <u>underline</u>: second best.

| Model | $\sigma$ | Selectivity ↑ | On-Target ↑ | Off-Target ↓ |
| --- | --- | --- | --- | --- |
| Native | 0 | 3.70% | <u>74.07%</u> | 74.07% |
| ProteinMPNN | 0.02 | <u>7.41%</u> | **77.78%** | 75.93% |
| ESM-IF | 0 | <u>7.41%</u> | **77.78%** | <u>72.22%</u> |
| PiFold | 0 | 3.70% | 72.22% | 75.93% |
| RedNet | 0.02 | 5.56% | <u>74.07%</u> | <u>72.22%</u> |
| RedNet-CD | 0.02 | **9.26%** | 72.22% | **70.37%** |

### 3.4 Structural analysis of redesigned selective binders

We illustrate how contrastive decoding enables selective redesign with two case studies (Figures 2 and 3).

**Figure 2:**
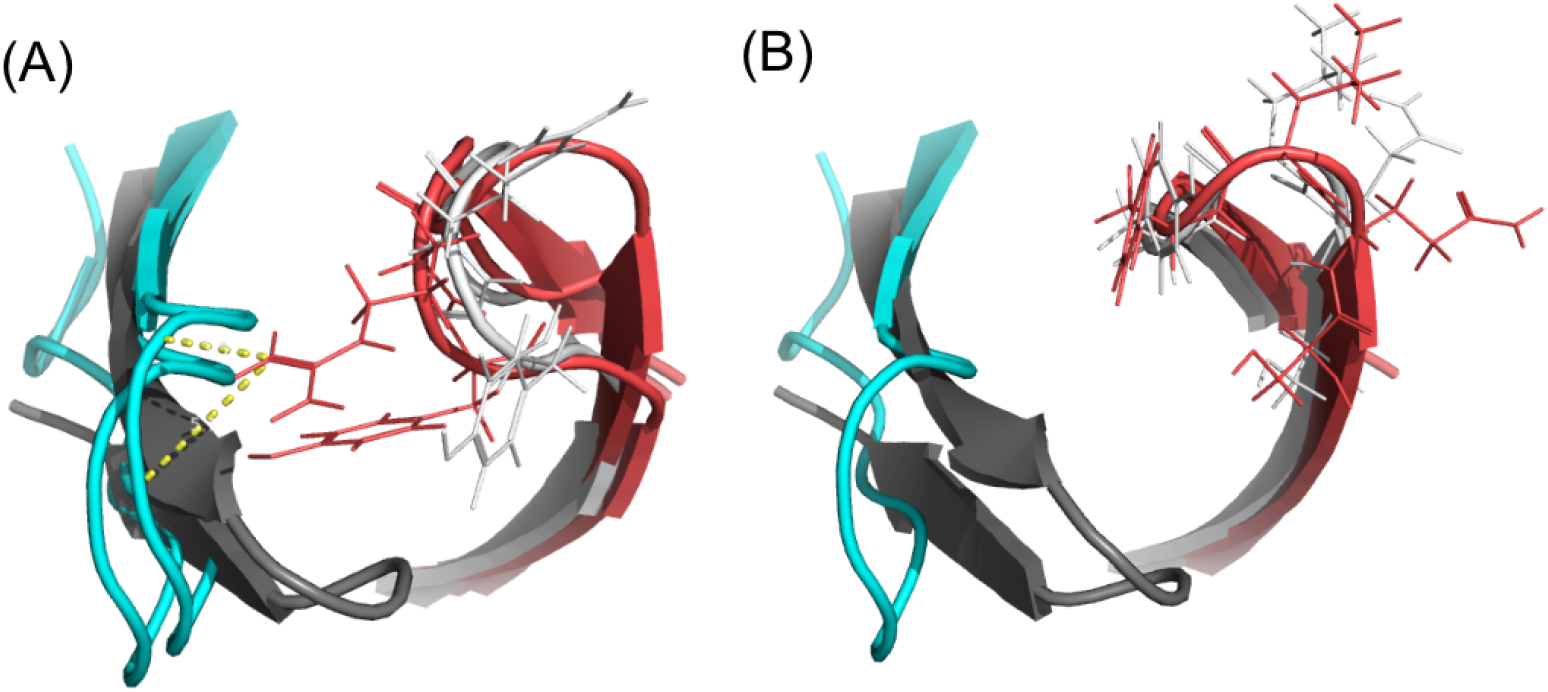
Structural analysis of the 6FOE–5WHJ selective binder pair. (A) Interactions of re-designed binders (red for the design chain of the on-target complex and white for the design chain of the off-target complex) to their respective on-target (cyan) and off-target (grey) partners. (B) Interactions of native binders to their respective on-target and off-target partners.

**Figure 3:**
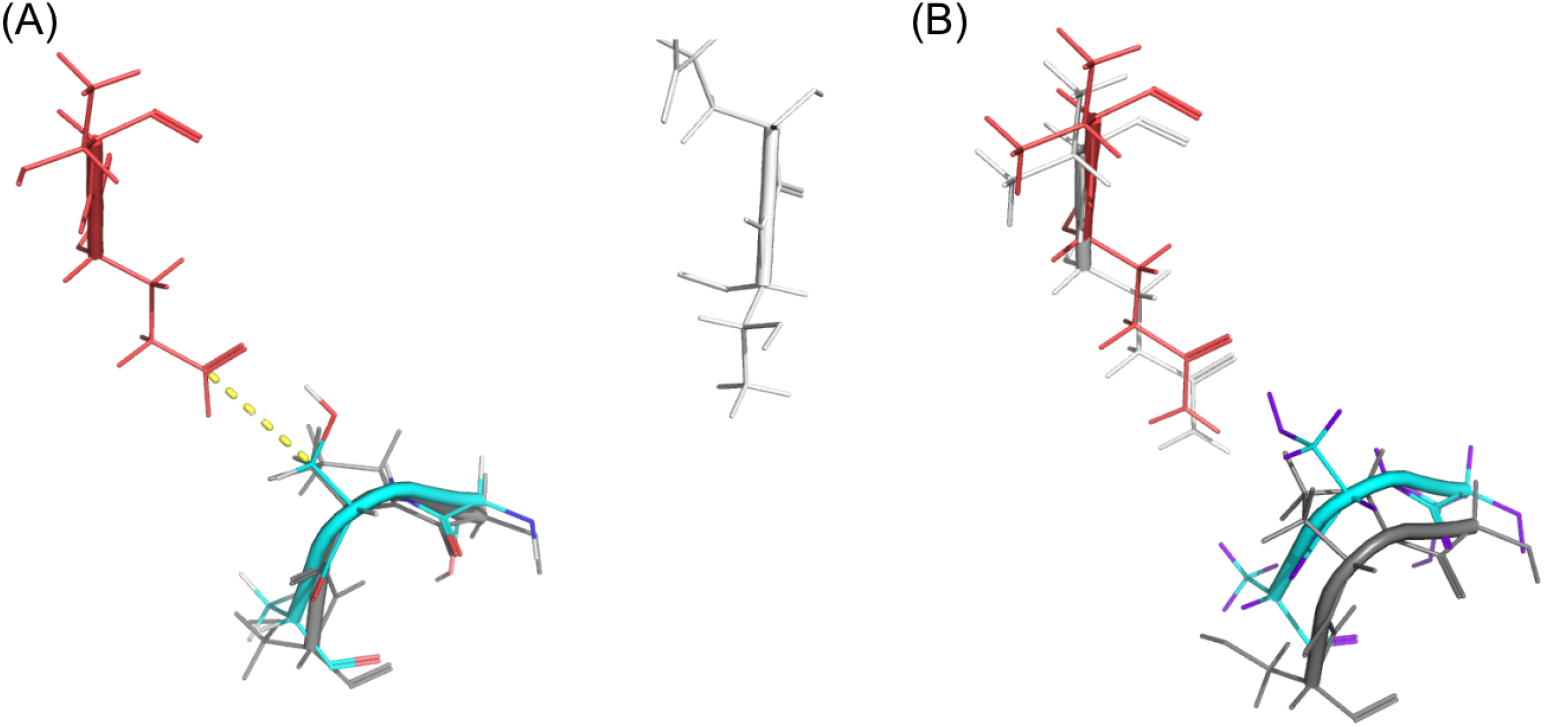
Structural analysis of the 5FFN–1LW6 selective binder pair. (A) Interactions of redesigned binders (red for the design chain of the on-target complex and white for the design chain of the off-target complex) to their respective on-target (cyan) and off-target (grey) partners. (B) Interactions of native binders to their respective on-target and off-target partners.

**Case 1: 6FOE–5WHJ (Fab)**. Both complexes are Fabs. The redesigned binder mutates four contiguous interface residues from SQLY to GYRN. The L →R substitution forms contacts with F and W on the 6FOE target but clashes with the corresponding 5WHJ residues (Figure 2A). Target backbones are nearly identical (RMSD 1.89 Å), so the selectivity comes entirely from side-chain differences.

**Case 2: 5FFN–1LW6 (Subtilisin)**. 5FFN and 1LW6 share the same target fold (Subtilisin), and their native binders are the chymotrypsin inhibitors CI2A and CI2. Target chains are structurally similar (RMSD 2.08 Å) and interfaces overlap substantially (Jaccard 0.38). The redesigned binder changes a QV segment to ET: ET retains strong interactions with the polar SSA patch on 5FFN but interacts poorly with the corresponding region of 1LW6 (Figure 3A). Selectivity here comes from destabilizing the off-target interface rather than enhancing the on-target one.

## 4 Conclusion

We presented RedNet, a fixed-backbone binder design framework combining an all-atom graph transformer with a contrastive decoding algorithm derived from the thermodynamic decomposition of binding free energy. Incorporating target side-chain geometry raises native sequence recovery on heterodimeric interfaces, and contrastive decoding matches native-sequence AlphaFold3 co-folding success on high-confidence targets while producing binding energetics, hydrogen bonding, and hydrophobicity on par with native interfaces. On a new selective-binder benchmark, contrastive decoding roughly doubles the base model’s energetic selectivity by exploiting side-chain differences between structurally similar on- and off-target interfaces. The decoding framework extends naturally to other multistate design tasks without retraining. The main limitation of the present work is the lack of wet-lab validation; we plan to address this in future work.

## References

[1] Deniz Akpinaroglu, Kosuke Seki, Amy Guo, Eleanor Zhu, Mark JS Kelly, and Tanja Kortemme. Structure-conditioned masked language models for protein sequence design generalize beyond the native sequence space. BioRxiv, pages 2023–12, 2023.

[2] Rebecca F. Alford, Andrew Leaver-Fay, Matthew Jeliazko R. Jeliazkov, J. O’Meara, Frank P. DiMaio, Hahnbeom Park, Maxim V. Shapovalov, P. Douglas Renfrew, Vikram K. Mulligan, Kalli Kappel, et al. The Rosetta all-atom energy function for macromolecular modeling and design. Journal of Chemical Theory and Computation, 13(6):3031–3048, 2017.

[3] Shaked Brody, Uri Alon, and Eran Yahav. How attentive are graph attention networks? In International Conference on Learning Representations (ICLR), 2022.

[4] Alexander E Chu, Tianyu Lu, and Po-Ssu Huang. Sparks of function by de novo protein design. Nature biotechnology, 42(2):203–215, 2024.

[5] Justas Dauparas, Ivan Anishchenko, Nathaniel Bennett, et al. Robust deep learning-based protein sequence design using proteinmpnn. Science, 378(6615):49–56, 2022.

[6] Henry Dieckhaus, Michael Brocidiacono, Nicholas Z Randolph, and Brian Kuhlman. Transfer learning to leverage larger datasets for improved prediction of protein stability changes. Proceedings of the national academy of sciences, 121(6):e2314853121, 2024.

[7] Zhangyang Gao, Cheng Tan, and Stan Z. Li. Pifold: Toward effective and efficient protein inverse folding. In International Conference on Learning Representations, 2023.

[8] Michael K Gilson, James A Given, Bruce L Bush, and J Andrew McCammon. The statistical-thermodynamic basis for predicting binding affinities: a physical framework. Biophysical Journal, 72(3):1047–1069, 1997.

[9] Amy B. Guo, Lindsey A. Kidd, Andrew J. Borst, Istvan Redl, Go Ueda, Xinyu Li, Sophie Chang, Jorge A. Fallas, Tanja Kortemme, and David Baker. Deep learning–guided design of dynamic proteins. Science, 388(6749):eadr7094, 2025.

[10] Mark A Hallen, Jonathan W Martin, Adegoke Ojewole, Jonathan D Jou, Anna U Lowegard, Marcel S Frenkel, Pablo Gainza, Hunter M Nisonoff, Aditya Mukund, Siyu Wang, et al. OSPREY 3.0: open-source protein redesign for you, with powerful new features. Journal of Computational Chemistry, 39(30):2494–2507, 2018.

[11] Lin Hong and Tanja Kortemme. An integrative approach to protein sequence design through multiobjective optimization. PLoS Computational Biology, 20(7):e1011953, 2024.

[12] Chloe Hsu, Robert Verkuil, Jason Liu, Zeming Lin, Brian L. Hie, Tom Sercu, Adam Lerer, and Alexander Rives. Learning inverse folding from millions of predicted structures. bioRxiv, 2022.

[13] Erika L. Humphris and David J. Mandell. A Rosetta-based algorithm for multi-state design of proteins. Structure, 13(2):313–323, 2005.

[14] John Ingraham, Vikas Garg, Regina Barzilay, and Tommi Jaakkola. Generative models for graph-based protein design. Advances in Neural Information Processing Systems, 32, 2019.

[15] Tanja Kortemme. De novo protein design—from new structures to programmable functions. Cell, 187(18):4934–4953, 2024.

[16] Brian Kuhlman and Philip Bradley. Advances in protein structure prediction and design. Nature reviews molecular cell biology, 20(11):681–697, 2019.

[17] Xiang Lisa Li, Ari Holtzman, Daniel Fried, Percy Liang, Jason Eisner, Tatsunori Hashimoto, Luke Zettlemoyer, and Mike Lewis. Contrastive decoding: Open-ended text generation as optimization. In Proceedings of the 61st Annual Meeting of the Association for Computational Linguistics (Volume 1: Long Papers), pages 12286–12312, Toronto, Canada, July 2023. Association for Computational Linguistics.

[18] Sean O’Brien and Mike Lewis. Contrastive decoding improves reasoning in large language models. arXiv preprint 2309.09117, 2023.

[19] C. Nick Pace, Hailong Fu, Katrina Lee Fryar, John Landua, Saul R. Trevino, David Schell, Richard L. Thurlkill, Satoshi Imura, J. Martin Scholtz, Ketan Gajiwala, et al. Contribution of hydrogen bonds to protein stability. Protein Science, 23(5):652–661, 2014.

[20] Martin Pacesa, Lennart Nickel, Christian Schellhaas, Joseph Schmidt, Ekaterina Pyatova, Lucas Kissling, Patrick Barendse, Jagrity Choudhury, Srajan Kapoor, Ana Alcaraz-Serna, et al. One-shot design of functional protein binders with bindcraft. Nature, 646(8084):483–492, 2025.

[21] Hahnbeom Park, Philip Bradley, Per Greisen Jr., Yuan Liu, David Baker, and Frank DiMaio. Simultaneous optimization of biomolecular energy functions on features from small molecules and macromolecules. Journal of Chemical Theory and Computation, 12(12):6201–6212, 2016.

[22] Joseph L Watson, David Juergens, Nathaniel R Bennett, Brian L Trippe, Jason Yim, Helen E Eisenach, Woody Ahern, Andrew J Borst, Robert J Ragotte, Lukas F Milles, et al. De novo design of protein structure and function with rfdiffusion. Nature, 620(7976):1089–1100, 2023.

[23] Ziwei Xie and Jinbo Xu. Deep graph learning of inter-protein contacts. Bioinformatics, 38(4):947–953, 2022.

[24] Vinicius Zambaldi, David La, Alexander E. Chu, Harshnira Patani, Amy E. Danson, Tristan O. C. Kwan, Thomas Frerix, Rosalia G. Schneider, David Saxton, Ashok Thillaisundaram, Zachary Wu, Isabel Moraes, Oskar Lange, Eliseo Papa, Gabriella Stanton, Victor Martin, Sukhdeep Singh, Lai H. Wong, Russ Bates, Simon A. Kohl, Josh Abramson, Andrew W. Senior, Yilmaz Alguel, Mary Y. Wu, Irene M. Aspalter, Katie Bentley, David L. V. Bauer, Peter Cherepanov, Demis Hassabis, Pushmeet Kohli, Rob Fergus, and Jue Wang. De novo design of high-affinity protein binders with AlphaProteo. arXiv preprint 2409.08022, 2024.

[25] Jie Zhou, Chau Q Le, Yun Zhang, and James A Wells. A general approach for selection of epitope-directed binders to proteins. Proceedings of the National Academy of Sciences, 121(19):e2317307121, 2024.

