## Supplemental File for "Redesign selective protein binders using contrastive decoding"

Ziwei Xie[1] and Jinbo Xu*[1,*]

#### S1 Zero-shot Affinity Prediction

##### S1.1 Contrastive Scoring

We score designed sequences with six log-likelihood variants. `ll` and `ll_global` are the mean log-likelihoods of the binder and full complex sequence under the bound structure. `ll_mt` averages over mutated positions only. `ll_ref` subtracts the wild-type log-likelihood at each mutated position, measuring whether each mutation is preferred over the wild-type residue.

The contrastive variants `ll_cd` and `ll_cd_ref` subtract the unbound-binder log-likelihood from the bound version, following the thermodynamic decomposition of $\Delta G$. `ll_cd` averages over the full binder chain; `ll_cd_ref` restricts to mutated positions and references the wild-type. Concurrent work [6, 7, 5] reports similar contrastive scores and finds improved correlation with experimental stability.

$$\begin{aligned}
\texttt{ll} &= \frac{1}{N_{\texttt{binder}}} \sum_{i=1}^{N_{\texttt{binder}}} \ell_i(a_i) \\
\texttt{ll_global} &= \frac{1}{N_{\texttt{complex}}} \sum_{i=1}^{N_{\texttt{complex}}} \ell_i(a_i) \\
\texttt{ll_mt} &= \frac{1}{|\mathcal{M}|} \sum_{i \in \mathcal{M}} \ell_i(a_i) \\
\texttt{ll_ref} &= \frac{1}{|\mathcal{M}|} \sum_{i \in \mathcal{M}} \big(\ell_i(a_i) - \ell_i(a_i^{\text{wt}})\big) \\
\texttt{ll_cd} &= \frac{1}{N_{\texttt{binder}}} \sum_{i=1}^{N_{\texttt{binder}}} \ell_i(a_i) \\
&\quad - \frac{1}{N_{\texttt{binder}}} \sum_{i=1}^{N_{\texttt{binder}}} \ell_i^u(a_i) \\
\texttt{ll_cd_ref} &= \frac{1}{|\mathcal{M}|} \sum_{i \in \mathcal{M}} \big(\ell_i(a_i) - \ell_i(a_i^{\text{wt}})\big) \\
&\quad - \frac{1}{|\mathcal{M}|} \sum_{i \in \mathcal{M}} \big(\ell_i^u(a_i) - \ell_i^u(a_i^{\text{wt}})\big)
\end{aligned}$$

Here $\ell_i = \log p_i$ is the predicted log-probability at position $i$ given the bound complex structure and the target sequence; $\ell_i^u$ is the same under the unbound binder structure; $a_i$ and $a_i^{\text{wt}}$ are the designed and wild-type residues at position $i$; $N_{\texttt{binder}}$ and $N_{\texttt{complex}}$ are the binder and complex sequence lengths; $\mathcal{M} = \{i : a_i \neq a_i^{\text{wt}}\}$ is the mutated-position set.

#### S1.2 Results

Table S1: Zero-shot binding affinity prediction on SKEMPI v2.0. Spearman correlations ($\rho$) averaged across assays. Scoring methods: standard binder log-likelihood (`ll`), global complex log-likelihood (`global`), mutated-residue log-likelihood (`mt`), reference-normalized (`ref`), contrastive log-likelihood (`cd_ll`), and contrastive reference-normalized (`cd_ll_ref`). $\sigma$: backbone coordinate noise level.

| Model | $\sigma$ | ll | global | mt | ref | cd_ll | cd_ll_ref |
|---|---|---|---|---|---|---|---|
| ESM-IF | 0 | 0.16 | 0.24 | 0.20 | 0.20 | 0.15 | 0.15 |
| PiFold | 0 | 0.17 | -0.17 | -0.03 | -0.12 | 0.15 | 0.00 |
| ProteinMPNN | 0.02 | 0.17 | 0.17 | 0.23 | 0.26 | 0.10 | 0.24 |
| RedNet | 0 | 0.18 | 0.23 | 0.21 | 0.22 | 0.23 | 0.26 |
| RedNet | 0.02 | 0.21 | 0.26 | 0.22 | 0.24 | 0.26 | 0.28 |

Supplementary Table S2 reports the full results.

Table S2: Zero-shot binding affinity prediction on SKEMPI v2.0. Spearman correlations ($\rho$), Kendall's $\tau$, and NDCG scores averaged across assays in SKEMPI v2.0. Contrastive scoring methods include standard binder log-likelihood (`ll`), global complex log-likelihood (`global`), mutated-residue log-likelihood (`mt`), reference-normalized (`ref`), contrastive log-likelihood (`cd_ll`), and contrastive reference-normalized (`cd_ll_ref`). $\sigma$: backbone coordinate noise level.

| Model | $\sigma$ | Spearman $\rho$ | | | | | | Kendall $\tau$ | | | | | | NDCG | | | | | |
|---|---|---|---|---|---|---|---|---|---|---|---|---|---|---|---|---|---|---|---|
| | | ll | global | mt | ref | cd_ll | cd_ll_ref | ll | global | mt | ref | cd_ll | cd_ll_ref | ll | global | mt | ref | cd_ll | cd_ll_ref |
| ESM-IF | 0 | 0.16 | 0.24 | 0.20 | 0.20 | 0.15 | 0.15 | 0.12 | 0.18 | 0.14 | 0.14 | 0.15 | 0.10 | 0.77 | 0.79 | 0.79 | 0.78 | 0.77 | 0.77 |
| PiFold | 0 | 0.17 | -0.17 | -0.03 | -0.12 | 0.15 | 0.00 | 0.13 | -0.13 | -0.02 | -0.09 | 0.11 | 0.00 | 0.78 | 0.67 | 0.72 | 0.69 | 0.75 | 0.71 |
| ProteinMPNN | 0.02 | 0.17 | 0.17 | 0.23 | 0.26 | 0.10 | 0.24 | 0.12 | 0.12 | 0.17 | 0.18 | 0.07 | 0.18 | 0.78 | 0.78 | 0.80 | 0.80 | 0.76 | 0.78 |
| RedNet | 0 | 0.18 | 0.23 | 0.21 | 0.22 | 0.23 | 0.26 | 0.13 | 0.17 | 0.15 | 0.16 | 0.17 | 0.18 | 0.79 | 0.80 | 0.80 | 0.80 | 0.78 | 0.78 |
| RedNet | 0.02 | 0.21 | 0.26 | 0.22 | 0.24 | 0.26 | 0.28 | 0.15 | 0.19 | 0.16 | 0.18 | 0.19 | 0.20 | 0.79 | 0.80 | 0.81 | 0.81 | 0.79 | 0.78 |

Prior work has shown contrastive scoring helps zero-shot stability prediction with ProteinMPNN and ESM-IF, but has not tested it for affinity prediction in a binder-design setting. We benchmark RedNet, ProteinMPNN, ESM-IF, and PiFold on SKEMPI v2.0 (Supplementary Tables S1 and S2). PiFold and ESM-IF have no released models trained at noise above $0.1$ Å, so all comparisons use $\sigma \in \{0, 0.02\}$.

**Overall.** RedNet ($\sigma$=$0.02$) with `cd_ll` or `cd_ll_ref` leads on every metric: Spearman's $\rho = 0.28$ (`cd_ll_ref`), Kendall's $\tau = 0.20$ (`cd_ll_ref`), NDCG $= 0.81$ (`mt`, `ref`).

**Across models.** RedNet ($\sigma$=$0.02$) gives the highest Spearman's $\rho$ on `ll` (0.21 vs. 0.17 for ProteinMPNN and ESM-IF) and `global` (0.26 vs. ESM-IF 0.24, ProteinMPNN 0.17), and leads on Kendall's $\tau$ in five of six scoring methods. ProteinMPNN edges out RedNet on Spearman's $\rho$ for `mt` (0.23 vs. 0.22) and `ref` (0.26 vs. 0.24); RedNet matches or beats it on Kendall's $\tau$ and NDCG for those metrics.

**Contrastive scoring is selective.** It improves RedNet at both noise levels on every metric, but degrades ProteinMPNN ($\rho$ drops from 0.26 with `ref` to 0.24 with `cd_ll_ref`; from 0.23 with `mt` to

Table S3: Architecture ablation of RedNet on monomers, homodimers, and heterodimers at backbone coordinate noise level $\sigma = 0.02$, evaluated on the same low-homology PDB test split as the main benchmark. NSR: native sequence recovery. LL: log-likelihood. PPL: perplexity. *Base* is the default RedNet configuration: a residue-graph encoder/decoder where each layer combines a mean-pool message-passing branch and a graph-attention (GAT) branch, with edge features built from C$\alpha$–C$\alpha$ RBF distances, frame shifts, and CB-to-side-chain RBF distances. The remaining rows modify a single component of *Base*. *w/o GAT*: drops the GAT branch from every encoder and decoder layer, leaving only the mean-pool message-passing path. *w/o side-chain*: drops the CB-to-side-chain RBF distance feature from edge representations; the kNN graph topology, frame shifts, and C$\alpha$–C$\alpha$ distances are unchanged. + *FA encoder*: adds an EGNN-style equivariant graph-attention layer to the featurizer that aggregates all atoms within a 10 Å centroid-to-atom radius graph into per-residue node features. + *FA-SE(3)*: extends + *FA encoder* with an auxiliary head on the same equivariant layer that predicts SE(3)-equivariant updates to the four backbone atom positions (N, C$\alpha$, C, O), supervised by an SVD-aligned MSE loss. + *edge loss*: adds an auxiliary edge-pair cross-entropy prediction head and loss to *Base*..

| Variant | Monomer | | | Homodimer | | | Heterodimer | | |
|---|---|---|---|---|---|---|---|---|---|
| | NSR | LL | PPL | NSR | LL | PPL | NSR | LL | PPL |
| Base | 0.37 | -1.9 | 8.3 | 0.44 | -1.4 | 4.6 | 0.39 | -1.9 | 7.5 |
| w/o GAT | 0.37 | -1.9 | 8.6 | 0.43 | -1.5 | 4.7 | 0.37 | -2.0 | 8.3 |
| w/o side-chain | 0.37 | -1.9 | 7.8 | 0.44 | -1.4 | 4.6 | 0.37 | -2.0 | 8.1 |
| + FA encoder | 0.37 | -1.9 | 9.4 | 0.44 | -1.4 | 4.5 | 0.39 | -1.9 | 7.9 |
| + FA-SE(3) | 0.38 | -1.9 | 8.6 | 0.44 | -1.4 | 4.5 | 0.38 | -2.0 | 8.1 |
| + edge loss | 0.37 | -1.9 | 8.2 | 0.43 | -1.5 | 4.7 | 0.39 | -1.9 | 7.7 |

0.10 with `cd_ll`) and ESM-IF ($\rho$ drops from 0.24 with `global` to 0.15 with `cd_ll` or `cd_ll_ref`). PiFold's baseline is near zero ($\rho = -0.03$ on `mt`, $-0.12$ on `ref`), so its contrastive-scoring gain is uninformative. RedNet trains on both bound complexes and monomers and uses all-atom interface features, which sharpens its bound-vs-unbound likelihood gap.

**Noise augmentation.** RedNet at $\sigma$=0.02 beats $\sigma$=0 on every scoring method: $\rho$ on `ll` rises from 0.18 to 0.21, on `global` from 0.23 to 0.26, on `cd_ll` from 0.23 to 0.26, on `cd_ll_ref` from 0.26 to 0.28. Backbone-coordinate noise sharpens the model's response to mutations, even though it lowers native sequence recovery.

All-atom training, noise augmentation, and contrastive scoring together give the strongest zero-shot affinity ranking, and motivate a decoder that exploits the same contrastive signal.

### S2 Ablation Studies

#### S2.1 Architecture ablation on sequence recovery

Supplementary Table S3 compares ablation variants. NSR differences across variants stay within 0.02 on every interface type. *Base* ties or leads every column and gives the lowest heterodimer perplexity. The all-atom variants (+ *FA encoder* and + *FA-SE(3)*) match or slightly trail *Base* on heterodimers; the residue graph with side-chain RBF distances already encodes the relevant interface geometry. Removing the GAT branch or the side-chain RBF feature drops heterodimer NSR and raises perplexity. The edge loss is neutral on NSR.

##### S2.2 Contrastive-decoding hyperparameters on the heterodimer co-folding benchmark

Table S4: Contrastive-decoding hyperparameter ablation on the heterodimer co-folding benchmark (107 targets; high-confidence subset of 44 targets where native AlphaFold3 predictions exceed pLDDT 70). Top block: AF3 co-folding metrics (success rate uses BindCraft criteria pLDDT$>80$, pTM$>0.55$, ipTM$>0.5$). Bottom block: Rosetta energetics on the intersected 91-target set after three relax repeats. We report both median and mean for binder_score because the mean is sensitive to a handful of targets where Rosetta returns very large negative scores under base/no-cd but not under the other CD settings; the median ranks the variants the same way with smaller magnitudes. *Base* is $\alpha$=1, $\beta$=0.9..

| Variant | All targets ($n$=107) | | | | High-confidence ($n$=44) | | | |
|---|---|---|---|---|---|---|---|---|
| | ranking | pTM | ipTM | success | ranking | pTM | ipTM | success |
| no-cd (control) | 0.49 | 0.54 | 0.37 | 0.25 | 0.70 | 0.76 | 0.56 | 0.57 |
| no-cd, edge loss | 0.48 | 0.53 | 0.37 | 0.31 | 0.68 | 0.75 | 0.55 | 0.59 |
| $\alpha$=1, $\beta$=0.9 (base) | 0.49 | 0.53 | 0.37 | 0.30 | 0.69 | 0.75 | 0.57 | 0.68 |
| $\alpha$=2, $\beta$=0.9 | 0.50 | 0.54 | 0.38 | 0.30 | 0.69 | 0.75 | 0.56 | 0.64 |
| $\alpha$=0.5, $\beta$=0.9 | 0.48 | 0.53 | 0.36 | 0.26 | 0.69 | 0.75 | 0.56 | 0.57 |
| $\alpha$=1, $\beta$=0.7 | 0.49 | 0.54 | 0.37 | 0.28 | 0.70 | 0.76 | 0.57 | 0.59 |
| $\alpha$=1, $\beta$=0.5 | 0.49 | 0.53 | 0.36 | 0.25 | 0.69 | 0.76 | 0.56 | 0.57 |
| $\alpha$=1, $\beta$=0.3 | 0.49 | 0.53 | 0.37 | 0.28 | 0.70 | 0.76 | 0.57 | 0.64 |

*Rosetta energetics on intersection ($n$=91, three replicas)*

| Variant | Binding | | | | Interface | | | |
|---|---|---|---|---|---|---|---|---|
| | binder_score (med) | binder_score (mean) | $\Delta G$ | dG/SASA | sc | packstat | nres | hbonds |
| no-cd (control) | −40.80 | −179.88 | −52.49 | −2.77 | 0.66 | 0.54 | 19.4 | 6.21 |
| no-cd, edge loss | −32.79 | −137.82 | −52.58 | −2.81 | 0.67 | 0.53 | 19.1 | 5.99 |
| $\alpha$=1, $\beta$=0.9 (base) | −42.85 | −182.26 | −54.47 | −2.78 | 0.66 | 0.54 | 19.7 | 7.01 |
| $\alpha$=2, $\beta$=0.9 | −34.14 | −139.07 | −53.93 | −2.75 | 0.65 | 0.55 | 19.3 | 6.80 |
| $\alpha$=0.5, $\beta$=0.9 | −35.18 | −136.86 | −50.55 | −2.68 | 0.66 | 0.52 | 18.2 | 6.16 |
| $\alpha$=1, $\beta$=0.7 | −35.40 | −136.25 | −51.26 | −2.65 | 0.64 | 0.54 | 18.8 | 5.90 |
| $\alpha$=1, $\beta$=0.5 | −32.89 | −135.44 | −55.14 | −2.68 | 0.65 | 0.54 | 20.0 | 6.26 |
| $\alpha$=1, $\beta$=0.3 | −32.85 | −133.24 | −52.63 | −2.76 | 0.65 | 0.54 | 18.4 | 6.47 |

We sweep contrastive strength $\alpha$ and truncation threshold $\beta$ on the 107-target heterodimer benchmark and report AlphaFold3 cofold metrics and Rosetta energetics (Supplementary Table S4). The base setting $\alpha$=1, $\beta$=0.9 gives the highest cofold success on the high-confidence subset (0.68) and the lowest Rosetta binding score ($-182.26$ REU). $\alpha$=2 and $\alpha$=0.5 both lower cofold success with little energy change. Tightening $\beta$ to 0.7, 0.5, or 0.3 shrinks the candidate set and worsens binding score (down to $-133.24$ REU) and interface hydrogen-bond count; pTM and ipTM stay roughly flat. The no-cd control (edge loss only) trails the base CD setting on energy, so the energy gain comes from the decoding rule, not the auxiliary loss.

##### S2.3 Contrastive-decoding hyperparameters on the selective-binder benchmark

We repeat the $(\alpha, \beta)$ sweep on the selective-binder benchmark (Supplementary Table S5), restricted to the confident-pair subset (Jaccard $< 0.5$, both targets with native pLDDT $> 70$; $n$=54). The base setting leads at the low (Diff $< 0$: 0.65 vs. 0.33 for no-cd) and medium (Diff $< -5$: 0.52) thresholds. $\alpha$=2 wins at the strict threshold (Diff $< -10$: 0.39), trading low-threshold selectivity for tighter energy gaps. Lowering $\beta$ to 0.7, 0.5, or 0.3 hurts iptm sel-succ but keeps energetic selectivity competitive.

Table S5: Contrastive-decoding hyperparameter ablation on the selective-binder benchmark. iptm sel-succ requires on-target ipTM $> 0.55$ and off-target ipTM $< 0.55$; energy sel-succ at threshold $\tau$ requires binder_score$_{\text{on}}$ − binder_score$_{\text{off}}$ $< \tau$. Restricted to the difficulty subset (Jaccard $< 0.5$) of confident pairs (both on/off receptors with native target-chain pLDDT $> 70$); $n$=54. *Base* is $\alpha$=1, $\beta$=0.9..

| Variant | ipTM sel-succ | Energy sel-succ $< 0$ | Energy sel-succ $< -5$ | Energy sel-succ $< -10$ | ipTM diff |
|---|---|---|---|---|---|
| no-cd (control) | 0.06 | 0.33 | 0.28 | 0.22 | +0.01 |
| $\alpha$=1, $\beta$=0.9 (base) | 0.09 | 0.65 | 0.52 | 0.33 | +0.02 |
| $\alpha$=2, $\beta$=0.9 | 0.07 | 0.63 | 0.48 | 0.39 | +0.01 |
| $\alpha$=0.5, $\beta$=0.9 | 0.04 | 0.50 | 0.37 | 0.28 | +0.01 |
| $\alpha$=1, $\beta$=0.7 | 0.02 | 0.59 | 0.44 | 0.35 | −0.00 |
| $\alpha$=1, $\beta$=0.5 | 0.06 | 0.46 | 0.44 | 0.31 | −0.02 |
| $\alpha$=1, $\beta$=0.3 | 0.07 | 0.52 | 0.44 | 0.31 | +0.01 |

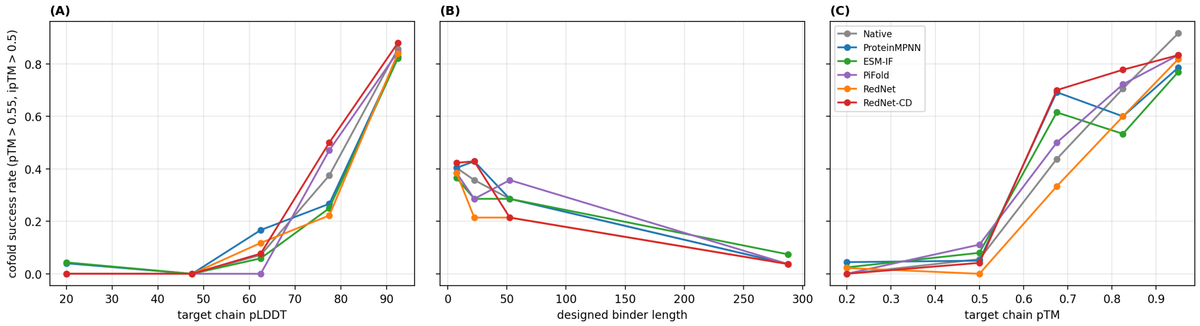


Figure S1: Factor analysis on the 107-target heterodimer benchmark. Cofold success = (pTM$>0.55$) $\wedge$ (ipTM$>0.5$). Each line is one method (labels match the heterodimer energetics table in the main text); each bin reports the mean success rate over (method, target) rows. (A) vs target-chain pLDDT (the pLDDT = 70 cutoff used for the high-confidence subset is between the third and fourth bin). (B) vs designed binder length. (C) vs target-chain pTM.

### S3 Factor Analysis

#### S3.1 Heterodimer co-folding

Supplementary Figure S1 stratifies cofold success on the 107-target benchmark by three target features. Target-chain confidence (pLDDT in A, pTM in C) drives the predicted confidence of the designed interface: success stays near zero when the target is poorly folded and rises sharply once the target is confident. The pLDDT $70$ cutoff used for the main-text high-confidence subset falls between the third and fourth pLDDT bin, on the inflection. Binder length (B) is anti-correlated with success: short binders ($< 30$ residues) succeed most often, and the rate drops as length grows.

#### S3.2 Selective binder

Supplementary Figure S2 stratifies energetic selectivity (binder_score$_{\text{on}}$ − binder_score$_{\text{off}}$ $< 0$) by three (on, off) features: Jaccard interface overlap, receptor sequence identity, and binder length. We omit the ipTM-based criterion because it is too sparse to discriminate methods at this scale (mean $7\%$ across methods). None of the three features shows a clear monotone trend with selectivity rate, so the across-method differences reported in the main text are not explained by any single benchmark-difficulty axis.

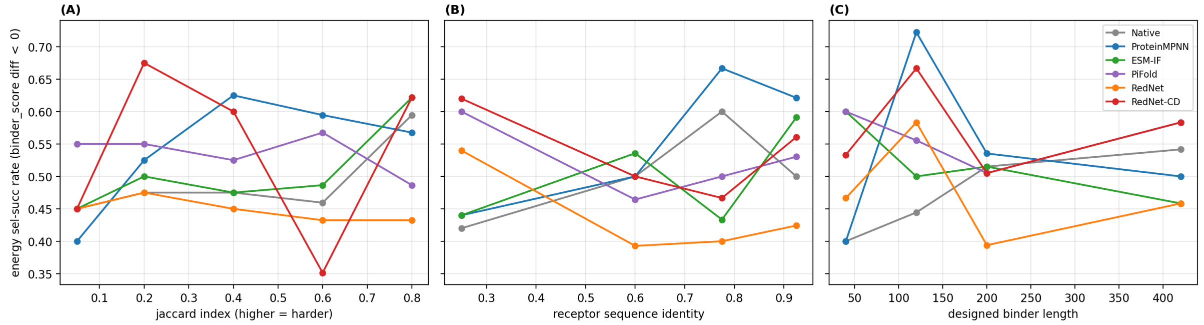


Figure S2: Factor analysis on the selective-binder benchmark. Energy sel-succ requires $\text{binder\_score}_{\text{on}} - \text{binder\_score}_{\text{off}} < 0$. Each line is one method (labels match Table S5); each bin reports the mean rate over (method, on/off pair) rows ($n{=}174$ pairs). (A) vs Jaccard index of inter-chain contact pairs (higher overlap = harder). (B) vs receptor sequence identity. (C) vs designed binder length. We omit the ipTM-based criterion because it is too sparse to discriminate methods at this scale (mean $7\%$ in Table S5).

### S4 Datasets

#### S4.1 Training Set

We use PDB [2] structures released before 2023-01-01. We discard structures with more than 20 polymer chains, resolution worse than $5\,\text{Å}$, or experimental methods other than X-ray diffraction and electron microscopy. We keep polypeptide(L) chains of 10–5000 residues with fewer than $10\%$ unknown residues.

**Validation set.** Structures released between 2022-05-01 and 2022-12-31 form the validation set, with redundant chains removed by the same procedure used for the test sets below.

#### S4.2 Test Datasets

**Low-homology PDB test set.** We start from structures released 2023-01-01 to 2023-12-31. To avoid sequence leakage, we MMseqs2-search [16] test chains against training and keep only design chains with e-value $> 1$, which is stricter than the common $30\%$ identity cutoff. We then cluster surviving sequences at $40\%$ identity. Each structure is labeled monomer (no interface), homodimer (similar interacting chains), or heterodimer (dissimilar). From the cluster representatives in each class we sample 300 monomers, 150 homodimers, and 150 heterodimers, for a 600-structure balanced test set.

For heterodimer co-folding we keep heterodimers with $\leq 500$ total residues for tractability, retaining 107 samples.

**Selective binder test set.** We curate heterodimeric complexes from PDB entries released before 2025-04-14. Two chains form an interacting heterodimer if their minimum $\text{C}\alpha$–$\text{C}\alpha$ distance within a bioassembly is $\leq 10\,\text{Å}$ and they share $< 90\%$ sequence identity (MMseqs2 clustering).

We keep chains of 20–500 residues with $< 10\%$ unknown residues and no single amino acid above $50\%$ frequency. For each chain cluster, we collect all interacting heterodimers and keep clusters with partners in at least 2 and at most 30 PDB entries; the upper bound excludes promiscuous clusters from dominating the set. This step yields 991 clusters and 3,246 heterodimers.

For each cluster (the binder cluster), we randomly pick one heterodimer as the on-target. Remaining heterodimers in the cluster are off-target candidates. We TM-align the binder chains (within-cluster) and the target chains separately. An off-target is dropped if (1) the target chains share $100\%$ identity, (2) binder alignment coverage is $< 90\%$, (3) binder alignment identity is $< 90\%$, or (4) binder alignment RMSD is $> 2.5\,\text{Å}$. After filtering, 691 clusters survive.

Off-target difficulty is the Jaccard similarity of inter-chain residue contact pairs ($\leq 10\,\text{Å}$ C$\alpha$–C$\alpha$):

$$\text{Difficulty} = \text{JaccardSimilarity}(\mathcal{C}_{\text{on}}, \mathcal{C}_{\text{off}})$$

where $\mathcal{C}_{\text{on}}$ and $\mathcal{C}_{\text{off}}$ are the inter-chain contact-pair sets of the on- and off-target complexes. Higher Jaccard means more shared interface contacts and harder selective design.

We keep pairs with Difficulty $< 0.9$ (656 clusters). Within each cluster, the most dissimilar off-target is chosen, giving 656 non-redundant on-/off-target pairs. Each pair requires two AlphaFold3 cofolding runs (10 recycles, 5 diffusion samples) and three Rosetta relaxations per complex. We uniformly sample 180 pairs for the benchmark.

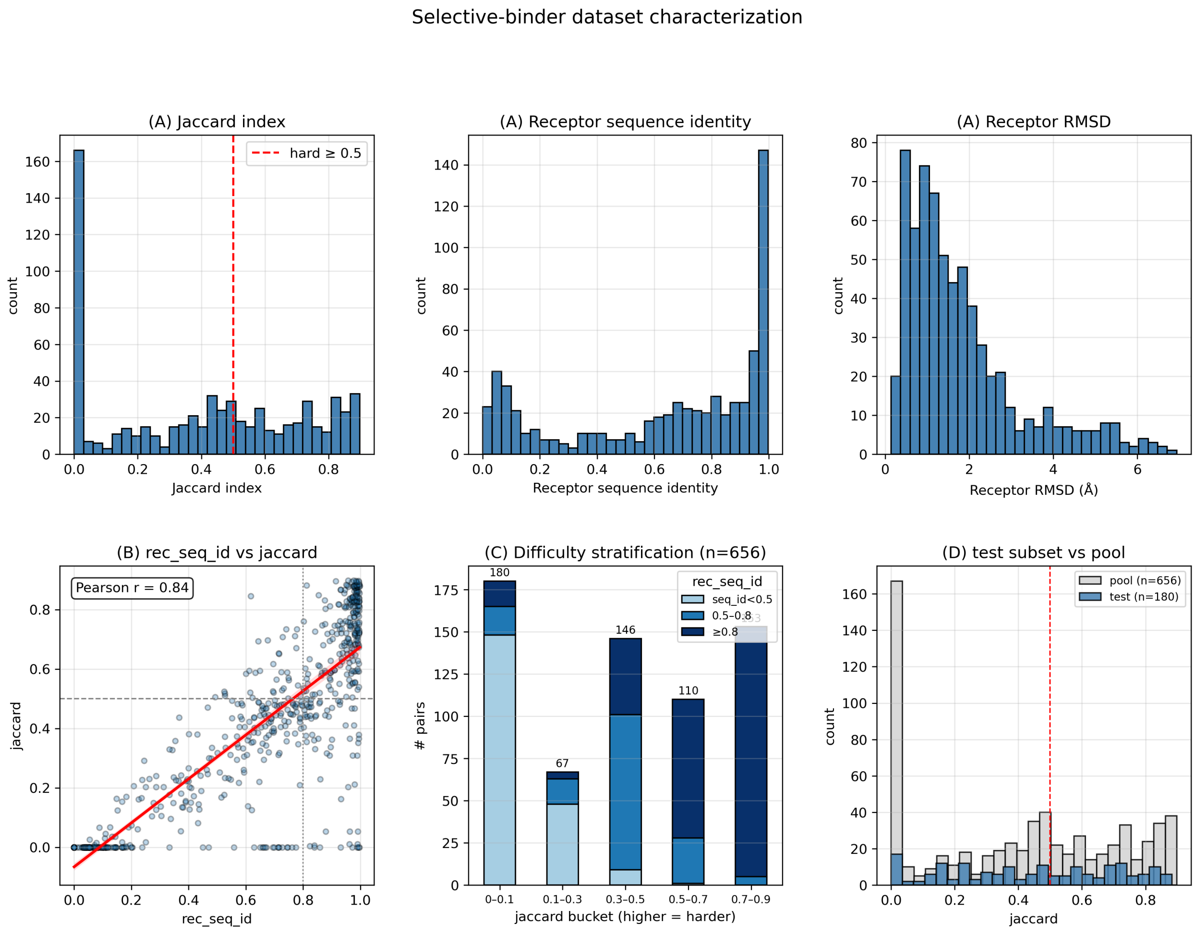


Figure S3: Selective-binder dataset. 656 pairs after cluster-level filtering; 180 sampled for evaluation. (A) Marginal distributions of difficulty features (Jaccard, receptor identity, receptor RMSD/TM-score, ligand identity/RMSD). (B) Receptor identity vs Jaccard, with linear fit (Pearson $r$=0.84). (C) 656-pair pool partitioned by Jaccard bucket and shaded by receptor identity. (D) 180-pair evaluation subset vs 656-pair pool; medians match within $0.03$, hard fraction $44\%$ vs $40\%$.

**Selective binder test set: difficulty profile.** Supplementary Figure S3 characterizes the dataset along two axes: Jaccard interface overlap and receptor sequence identity. The higher either score, the harder it is to distinguish the on- and off-target interactions. The 656-pair pool covers Jaccard from $0$ to $0.90$ (median $0.44$) and receptor identity from $0$ to $1.0$ (median $0.74$). The two axes correlate strongly (Pearson $r$=0.84) because homologous receptors share binding-site residues. 40% of pairs sit at Jaccard $\geq 0.5$ and 35% are hard on both axes simultaneously. The 180-pair evaluation subset preserves this profile (medians within 0.03, hard fraction 44%).

### S5 Feature Tables

Supplementary Table S6 lists the node and edge features for the residue-level and atom-level graphs.

Table S6: Summary of input features. Core atoms: N, C$\alpha$, C, O, pseudo-C$\beta$ ($C$=5). The residue graph is a $K$-NN graph ($K$=48) over C$\alpha$ distances. The atom graph connects each C$\alpha$ to nearby atoms via a radius graph ($r$=15 Å, max $k$=96). RBF: $\phi(d) = \exp(-(d - \mu_i)^2)$, $\mu_i$ linearly spaced in $[2, 22]$, $D$=16 bins. Atom type vocabulary $A$=37; residue type vocabulary $R$=33. $N$: residues; $M$: atoms; $E$: atom graph edges.

| Feature | Shape | Description |
|---|---|---|
| *Residue graph: edge features* | | |
| Core atom pairwise RBF | $(N, K, C^2D)$ | RBF over pairwise distances between core atoms |
| Core atom inverse distance | $(N, K, C^2)$ | $(1 + d_{ij})^{-1}$ for each core atom pair |
| Relative residue index | $(N, K)$ | Sequence offset encoding |
| Same-chain indicator | $(N, K)$ | 1 if same chain |
| Frame-relative positions | $(N, K, 3C)$ | Core atom coordinates in local N–C$\alpha$–C frames |
| C$\beta$–sidechain RBF | $(N, K, 32D)$ | RBF from pseudo-C$\beta$ to sidechain atoms; design chains masked |
| *Atom graph: node features* | | |
| Atom type | $(M, A)$ | One-hot over atom types |
| Residue type | $(M, R)$ | One-hot over parent residue type |
| Atom exists | $(M, 1)$ | 1 if atom resolved in structure |
| *Atom graph: edge features* | | |
| RBF distance | $(E, D)$ | RBF over C$\alpha$-to-atom distance |
| Euclidean distance | $(E, 1)$ | C$\alpha$-to-atom distance |
| Residue index offset | $(E, 65)$ | Clamped relative residue index one-hot ($\pm$32) |
| Same-chain indicator | $(E, 1)$ | 1 if same chain |
| *Atom graph: 3D coordinates (equivariant attention)* | | |
| Centroid positions | $(N, 3)$ | C$\alpha$ coordinates |
| Atom positions | $(M, 3)$ | All-atom coordinates |

### S6 Hydrophobicity and Hydrogen Bonding of Designed Heterodimers

Supplementary Table S7 reports surface hydrophobicity and hydrogen bonding of designed interfaces, computed in Rosetta following BindCraft.

### S7 Architectures and Training

#### S7.1 All-atom equivariant GAT

For target side-chain modeling we use an SE(3)-invariant GAT (Supplementary Algorithm S2) that encodes messages from pairwise distances and edge attributes, with GATv2 attention selecting relevant neighboring atoms.

Table S7: Hydrophobicity and hydrogen-bond properties of designed interfaces. Surf Hydro (0–1): surface hydrophobicity. Int Nres: number of interface residues. Int HBonds: number of interface hydrogen bonds. Int HBond %: percentage of interface residues involved in hydrogen bonds. Int dUnsat HB: number of unsatisfied interface hydrogen bonds. Int dUnsat HB %: percentage of unsatisfied interface hydrogen bonds. Due to Rosetta relaxation failures, we analyze 91 of 107 heterodimers that are successfully relaxed for all methods. RedNet-Ens combines RedNet and RedNet-CD by selecting the design with the best binding score..

| Model | $\sigma$ | Surf Hydro ↓ | Int Nres ↑ | Int HBonds ↑ | Int HBond % ↑ | Int dUnsat HB ↓ | Int dUnsat HB % ↓ |
|---|---|---|---|---|---|---|---|
| Native | 0 | 0.43 | 20.10 | 6.97 | 46.15 | 2.98 | 17.90 |
| ProteinMPNN | 0.02 | 0.43 | 17.34 | 5.44 | 40.30 | 2.52 | 17.62 |
| ESM-IF | 0 | 0.46 | 19.92 | 6.51 | 43.19 | 2.78 | 17.26 |
| PiFold | 0 | 0.47 | 18.98 | 6.07 | 43.49 | 2.91 | 19.32 |
| RedNet | 0.02 | 0.45 | 19.42 | 6.21 | 44.80 | 2.44 | 15.40 |
| RedNet-CD | 0.02 | 0.44 | 19.70 | 7.01 | 48.23 | 2.63 | 14.77 |
| RedNet-Ens | 0.02 | 0.44 | 20.19 | 7.31 | 48.99 | 2.59 | 14.27 |

**Algorithm S1** GAT Layer

\# $s$: node features; $p$: edge features; $\mathcal{E}$: neighbor indices; $M$: edge mask
**def** GATLayer($s$, $p$, $\mathcal{E}$, $M$):

\# Build edge messages from nodes and edges
$m_{ij} \leftarrow \textsc{Linear}(p_{ij}) + \textsc{Gather}(W^{\text{src}} s, \mathcal{E}) + W^{\text{tgt}} s_i$
$m_{ij} \leftarrow \text{MLP}(m_{ij})$

\# Global pooling with gating
$o_i^{\text{global}} \leftarrow \frac{1}{|\delta(i)|} \sum_{j \in \delta(i)} m_{ij}$
$\Delta s \leftarrow \textsc{Linear}(\textsc{Sigmoid}(W^g s_i) \odot o_i^{\text{global}})$

\# Graph attention with gating
$\alpha_{ij} \leftarrow W^A \cdot \textsc{LeakyReLU}(\textsc{Linear}(m_{ij}))$
$\alpha_{ij} \leftarrow \textsc{Softmax}_{j \in \delta(i)}(\alpha_{ij})$
$v_{ij} \leftarrow \textsc{Linear}(m_{ij})$
$o_i^{\text{gat}} \leftarrow \sum_{j \in \delta(i)} \alpha_{ij} v_{ij}$
$\Delta s \leftarrow \Delta s + \textsc{Linear}(\textsc{Sigmoid}(W^{g'} s_i) \odot o_i^{\text{gat}})$

\# Residual updates
$s \leftarrow s + \text{Dropout}(\Delta s)$
$s \leftarrow s + \text{Dropout}(\text{MLP}(s))$
**return** $s$, $p$

#### S7.2 Pseudocode

Pseudocode for the GAT layer (Supplementary Algorithm S1) and the equivariant graph attention layer (Supplementary Algorithm S2).

#### S7.3 Losses

We train with a cross-entropy loss over amino acid tokens [13]. Given predicted logits $\hat{y}_i \in \mathbb{R}^{|\mathcal{V}|}$ and ground-truth token $y_i$ at position $i$, the nodewise loss is

$$\mathcal{L}_{\text{node}} = \sum_{m \in \mathcal{M}} w_m \cdot \frac{1}{|M_m|} \sum_{i \in M_m} \text{CE}(\hat{y}_i, y_i),$$

**Algorithm S2** Equivariant Graph Attention Layer

\# $q$, $k$: node features; $x$, $y$: coordinates; $\mathcal{E}$: edges; $e$: edge features
**def** EGATLAYER($q$, $k$, $x$, $y$, $\mathcal{E}$, $e$):

\# Compute relative positions and distances
$(i, j) \leftarrow \mathcal{E}$
$z_{ij} \leftarrow y_j - x_i$
$d_{ij} \leftarrow \|z_{ij}\|^2$

\# Build edge messages
$m_{ij} \leftarrow \textsc{Linear}([q_i \| k_j \| d_{ij} \| e_{ij}])$

\# Compute multi-head attention
$\alpha_{ij} \leftarrow W^Q q_i + W^K k_j + W^E m_{ij}$
$\alpha_{ij} \leftarrow W^A \cdot \textsc{LeakyReLU}(\alpha_{ij})$
$\alpha_{ij} \leftarrow \textsc{Softmax}_{j \in \delta(i)}(\alpha_{ij})$ # over neighbors $j$

\# Aggregate values
$v_{ij} \leftarrow \textsc{Linear}([q_i \| k_j \| d_{ij} \| e_{ij}])$
$o_i \leftarrow \sum_{j \in \delta(i)} \alpha_{ij} v_{ij}$
$h^{\text{out}} \leftarrow \textsc{Linear}([q_i \| o_i])$
**return** $h^{\text{out}}$

where $\mathcal{M}$ is a set of binary masks (e.g., design-site, prediction) with weights $w_m$ and CE$(\cdot, \cdot)$ is the cross-entropy loss.

We add an auxiliary edgewise loss that prevents overfitting. For each node $i$ and its $k$-nearest neighbors, the model predicts a joint token distribution over residue pairs. Given edgewise logits $\hat{z}_{ij} \in \mathbb{R}^{|\mathcal{V}|^2}$ and the pair token $z_{ij} = y_i \cdot |\mathcal{V}| + y_j$, $\mathcal{L}_{\text{edge}} = \frac{1}{|E|} \sum_{(i,j) \in E} \text{CE}(\hat{z}_{ij}, z_{ij})$, over valid edges $E$. The total loss is $\mathcal{L} = \mathcal{L}_{\text{node}} + \lambda_{\text{edge}} \mathcal{L}_{\text{edge}}$, with $\lambda_{\text{edge}} = 1$.

### S8 Related Works

**Fixed-backbone design.** Physics-based methods couple an energy function for van der Waals, hydrogen bonding, electrostatics, and solvation [1, 15] with a sequence-space search. Rosetta uses simulated annealing [1]; OSPREY uses dead-end elimination to find provably minimum-energy sequences [10]. Both have produced experimental successes but scale poorly and lean on experimental screening to identify functional designs.

**Deep learning for fixed-backbone design.** Deep learning has improved both efficiency and wet-lab success rates. Autoregressive models—Structured Transformer [13], ProteinMPNN [4]—decode sequences iteratively over a graph encoder. Non-autoregressive models such as PiFold [8] produce a full sequence in one pass at competitive accuracy. ProteinMPNN reports much higher experimental success than Rosetta on monomers [4] and is now standard in de novo design pipelines.

**One-sided interface design.** Designing a binder against a fixed target spans applications from receptor modulation to pathogen neutralization. Physics-based approaches succeed mostly on targets with hydrophobic patches or concave surfaces and on small stable scaffolds [3].

Deep learning has reshaped the field. RFdiffusion [17] generates binder backbones via diffusion, then sequences with ProteinMPNN. BindCraft [14] directly optimizes sequences through AlphaFold2 confidence backpropagation. Both produce validated binders, but per-target success rates span $0\%$ to $> 90\%$ [18] and many designs need further optimization.

**Multistate design for binding specificity.** Selective binding requires multistate optimization. Rosetta MSD [12] maximizes energy gaps across states. Deep-learning extensions add fixed-backbone scoring [11] or structural ensembles, with success on conformational switches [9]; their reach to general specificity tuning is unclear. Differential yeast display [19] identifies selective binders experimentally without using computational design directly.

### References


[1] Rebecca F. Alford, Andrew Leaver-Fay, Jeliazko R. Jeliazkov, Matthew J. O'Meara, Frank P. DiMaio, Hahnbeom Park, Maxim V. Shapovalov, P. Douglas Renfrew, Vikram K. Mulligan, Kalli Kappel, et al. The Rosetta all-atom energy function for macromolecular modeling and design. *Journal of Chemical Theory and Computation*, 13(6):3031–3048, 2017.

[2] Helen M Berman, John Westbrook, Zukang Feng, Gary Gilliland, T N Bhat, Helge Weissig, Ilya N Shindyalov, and Philip E Bourne. The protein data bank. *Nucleic acids research*, 28(1):235–242, 2000.

[3] Aaron Chevalier, Daniel-Adriano Silva, Gabriel J Rocklin, David R Hicks, Rozane Gebabla, et al. Massively parallel de novo protein design for targeted therapeutics. *Nature*, 550(7674):74–79, 2017.

[4] Justas Dauparas, Ivan Anishchenko, Nathaniel Bennett, et al. Robust deep learning-based protein sequence design using proteinmpnn. *Science*, 378(6615):49–56, 2022.

[5] Arthur Deng, Karsten D. Householder, Fang Wu, K. Christopher Garcia, and Brian L. Trippe. Predicting mutational effects on protein binding from folding energy. In *Proceedings of the 42nd International Conference on Machine Learning (ICML)*, volume 267 of *Proceedings of Machine Learning Research*, pages 13129–13151. PMLR, 2025.

[6] Oliver Dutton, Sandro Bottaro, Michele Invernizzi, et al. Improving inverse folding models at protein stability prediction without additional training or data. *Biophysics*, 2024.

[7] Jes Frellsen, M M Kassem, T Bengtsen, et al. Zero-shot protein stability prediction by inverse folding models: a free energy interpretation. *arXiv preprint arXiv:2506.05596*, 2025.

[8] Zhangyang Gao, Cheng Tan, and Stan Z. Li. Pifold: Toward effective and efficient protein inverse folding. In *International Conference on Learning Representations*, 2023.

[9] Amy B. Guo, Lindsey A. Kidd, Andrew J. Borst, Istvan Redl, Go Ueda, Xinyu Li, Sophie Chang, Jorge A. Fallas, Tanja Kortemme, and David Baker. Deep learning–guided design of dynamic proteins. *Science*, 388(6749):eadr7094, 2025.

[10] Mark A Hallen, Jonathan W Martin, Adegoke Ojewole, Jonathan D Jou, Anna U Lowegard, Marcel S Frenkel, Pablo Gainza, Hunter M Nisonoff, Aditya Mukund, Siyu Wang, et al. OSPREY 3.0: open-source protein redesign for you, with powerful new features. *Journal of Computational Chemistry*, 39(30):2494–2507, 2018.

[11] Lin Hong and Tanja Kortemme. An integrative approach to protein sequence design through multiobjective optimization. *PLoS Computational Biology*, 20(7):e1011953, 2024.

[12] Erika L. Humphris and David J. Mandell. A Rosetta-based algorithm for multi-state design of proteins. *Structure*, 13(2):313–323, 2005.

[13] John Ingraham, Vikas Garg, Regina Barzilay, and Tommi Jaakkola. Generative models for graph-based protein design. *Advances in Neural Information Processing Systems*, 32, 2019.

[14] Martin Pacesa, Lennart Nickel, Christian Schellhaas, Joseph Schmidt, Ekaterina Pyatova, Lucas Kissling, Patrick Barendse, Jagrity Choudhury, Srajan Kapoor, Ana Alcaraz-Serna, et al. One-shot design of functional protein binders with bindcraft. *Nature*, 646(8084):483–492, 2025.

[15] Hahnbeom Park, Philip Bradley, Per Greisen Jr., Yuan Liu, David Baker, and Frank DiMaio. Simultaneous optimization of biomolecular energy functions on features from small molecules and macromolecules. *Journal of Chemical Theory and Computation*, 12(12):6201–6212, 2016.

[16] Martin Steinegger and Johannes Söding. Mmseqs2 enables sensitive protein sequence searching for the analysis of massive data sets. *Nature Biotechnology*, 35(11):1026–1028, 2017.

[17] Joseph L Watson, David Juergens, Nathaniel R Bennett, Brian L Trippe, Jason Yim, Helen E Eisenach, Woody Ahern, Andrew J Borst, Robert J Ragotte, Lukas F Milles, et al. De novo design of protein structure and function with rfdiffusion. *Nature*, 620(7976):1089–1100, 2023.

[18] Vinicius Zambaldi, David La, Alexander E. Chu, Harshnira Patani, Amy E. Danson, Tristan O. C. Kwan, Thomas Frerix, Rosalia G. Schneider, David Saxton, Ashok Thillaisundaram, Zachary Wu, Isabel Moraes, Oskar Lange, Eliseo Papa, Gabriella Stanton, Victor Martin, Sukhdeep Singh, Lai H. Wong, Russ Bates, Simon A. Kohl, Josh Abramson, Andrew W. Senior, Yilmaz Alguel, Mary Y. Wu, Irene M. Aspalter, Katie Bentley, David L. V. Bauer, Peter Cherepanov, Demis Hassabis, Pushmeet Kohli, Rob Fergus, and Jue Wang. De novo design of high-affinity protein binders with AlphaProteo. *arXiv preprint arXiv:2409.08022*, 2024.

[19] Jie Zhou, Chau Q Le, Yun Zhang, and James A Wells. A general approach for selection of epitope-directed binders to proteins. *Proceedings of the National Academy of Sciences*, 121(19):e2317307121, 2024.
